# Thalamocortical spectral transmission relies on balanced input strengths

**DOI:** 10.1101/2020.09.20.305136

**Authors:** Matteo Saponati, Jordi Garcia-Ojalvo, Enrico Cataldo, Alberto Mazzoni

## Abstract

The thalamus is a key element of sensory transmission in the brain, as all sensory information is processed by the thalamus before reaching the cortex. The thalamus is known to gate and select sensory streams through a modulation of its internal activity in which spindle oscillations play a preponderant role, but the mechanism underlying this process is not completely understood. In particular, how do thalamocortical connections convey stimulus-driven information selectively over the background of thalamic internally generated activity (such as spindle oscillations)? Here we investigate this issue with a spiking network model of connectivity between thalamus and primary sensory cortex reproducing the local field potential of both areas. We found two features of the thalamocortical dynamics that filter out spindle oscillations: i) spindle oscillations are weaker in neurons projecting to the cortex, ii) the resonance dynamics of cortical networks selectively blocks frequency in the range encompassing spindle oscillations. This latter mechanism depends on the balance of the strength of thalamocortical connections toward excitatory and inhibitory neurons in the cortex. Our results pave the way toward an integrated understanding of the sensory streams traveling between the periphery and the cortex.

## 1 Introduction

During the past decades several key features of sensory processing in the primary sensory cortex have been discovered, but much less is known about sensory information processing and transmission in the thalamus [1]. The majority of sensory signals are conveyed by the thalamus to the cortex in the form of spiking patterns propagating along different pathways [2]. Thalamocortical relay neurons in the thalamus receive sensory inputs and in turn project them to particular areas of the cortex through thalamocortical synapses. For decades, the thalamus has been described as a relay station where little computation takes place. However, more recently, experimental and theoretical findings have shown a prominent role of the thalamus in both pre-processing of sensory stimuli [3] and modulation of cortical activity even in absence of external stimulation [4, 5, 6]. Here we investigate the interplay of these two functions of the thalamus, using a novel spiking network model of the two areas.

Several brain structures exhibit oscillatory activitiy at different timescales related to various cognitive functions such as memory consolidation [7], attention [8,9] and information transmission [10]. State-of-the-art techniques in systems neuroscience provides a detailed description of anatomical and electrophisiological structures of thalamic and cortical systems. However, a rigorous analysis of the role of thalamic and cortical oscillations in information transmission is missing. During sleep or anesthesia, slow-waves [0.1-1 Hz or 1-4 Hz] are present in both thalamic and cortical activities and seem to coherently organize the dynamics of both networks [11]. On the other hand, fast rhythms in different frequency bands characterize the activity of the thalamocortical system during states of awakeness or REM sleep. The thalamus also shows rhythmic activities in the [7-14 Hz], the so-called spindle oscillations, in particular during sleep. While common in the thalamus, these rhythms are hardly observed in primary cortical dynamics during awake states [12]. The joint thalamocortical system thus includes neural rhythms at different frequencies, which will have differential impact on the frequency bands that are known to carry different information in the cortex [40]. Therefore, the capability of the thalamocortical system to filter specific rhythms will determine its fundamental role in gating information flow throughout the brain. Do spindle oscillations in the thalamus have a role in transfering information to the cortex? Or do they act instead as an internal thalamic mechanism poorly related to sensory stimulation?

Information throughout the brain is mainly transferred by excitatory populations through synaptic connections between different areas. In the thalamus, links from sub-cortical areas to the cortex are driven by AMPAergic thalamocortical relay (TC) neurons. These excitatory neurons are surrounded by a shell of GABAergic reticular neurons (RE) which do not receive sensory input directly (see Figure 1A) and are supposed to modulate the information flow [13]. They receive afferents from TC neurons and send inhibitory inputs back to them, creating a closed-loop between thalamic population. Therefore, the activity of TC neurons does not convey a faithfull reproduction of sensory information, but rather a pre-processed version of them in function of internal states of the thalamus. Intrinsic features of thalamocortical feedforward connectivity are indeed crucial in shaping this information transmission. However, a complete understanding of anatomical and functional connectivity from TC neurons to the cortex is missing.

**Fig. 1:**
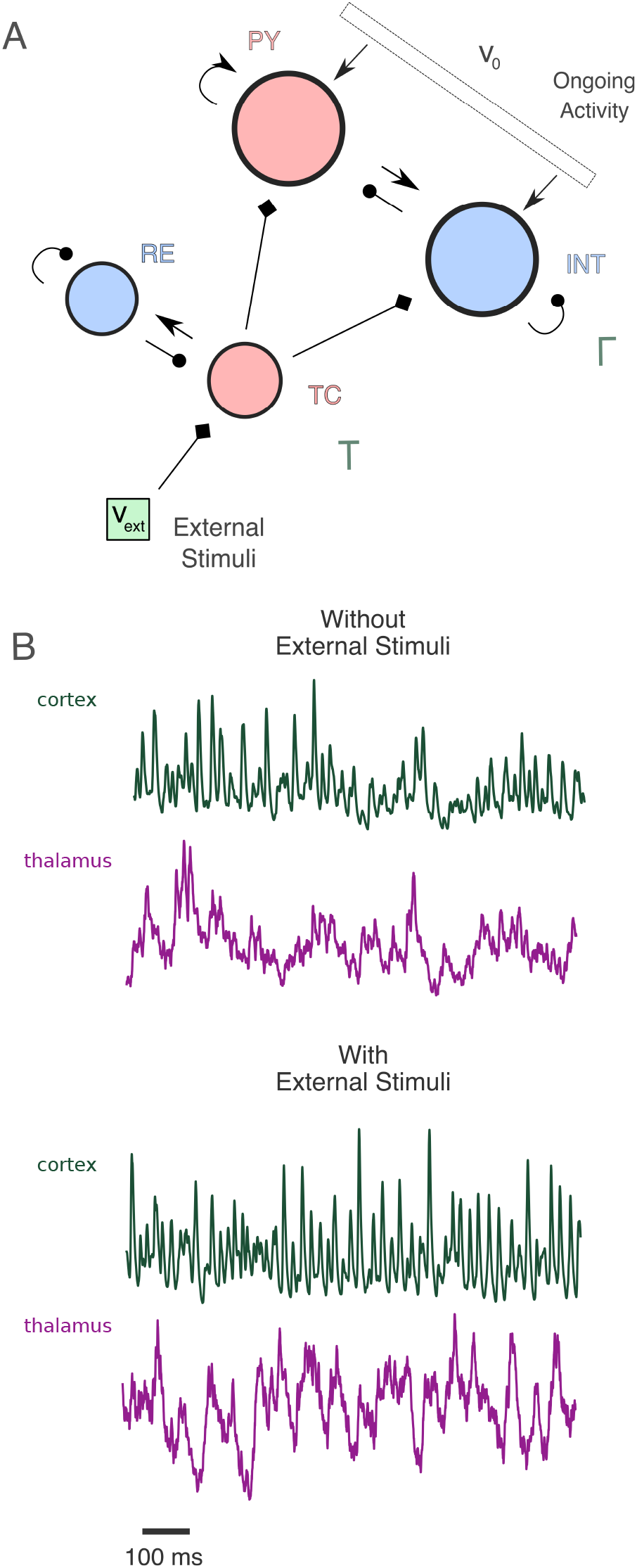
Network design and dynamics. A) Representation of the network structure. The thalamocortical model is composed by a thalamic network T and a cortical network Λ. Both networks are divided into an excitatory and an inhibitory population, in red and blue respectively. Arrows with triangle-shaped and circle-shaped heads represent excitatory and inhibitory connections, respectively. Arrows with diamond-shaped heads represent excitatory connections between different areas (External Stimuli to TC population, TC population to cortical populations). The cortical network receives background excitatory stimulation simulating ongoing activity of afferent cortical areas. B) Temporally aligned cortical LFP_T_(*t*) (green) and thalamic LFP_T_(*t*) (purple) from a numerical simulation in the absence (*ν*_ext_ = 0 spk/ms) or presence (*ν*_ext_ = 0.5 spk/ms) of external stimuli (see Methods).

In this scenario, we aim to shed new light on thalamic and cortical spindle oscillations basing our investigations on a novel integrate-and-fire network model [14] able to reproduce network oscillations on a wide range of timescales.

## 2 Methods

Here we summarize the main features of the model; we refer to [14] for further details. The network is an extension of a thalamic network model introduced in [34] extended to include a network model of a related cortical area following methods of [40,38].

### 2.1 Neural and Network Models

The thalamocortical network model consists of two structures, namely a thalamic network T and a cortical network Γ, see Figure 1A for a graphical representation of network structure. Both networks are composed by an excitatory and an inhibitory population. The thalamic network T is composed by 250 *thalamocortical relay* (TC) neurons with AMPA-like synapses and 250 *reticular* (RE) neurons with GABA-like synapses. The cortical network Γ is composed by 4000 *pyramidal* (PY) neurons with AMPA-like synapses and 1000 *interneurons* (INT) with GABA-like synapses. Thalamic and cortical structures are characterized by random and sparse connectivity schemes with different coupling probabilities. Any directed pair of cortical neurons are connected with a probability *p* = 0.02 independently from the neuron type, while thalamic RE neurons connect with TC neurons with a probability *p* = 0.04 and TC neurons connect to RE neurons with *p* = 0.01. Moreover, RE network structure shows recurrent connections with probability *p* = 0.04 on which we add a degree of clustering by starting from a ring network and then randomly rewiring with probability 0.25. This is necessary for the onset of sustained thalamic oscillations [34]. The model includes thalamocortical afferents by considering synaptic connections from TC neurons to cortical excitatory and inhibitory populations. Thalamocortical connectivity is random and sparse with a connection probability *p* = 0.07.

Neuronal dynamics is simulated with the Adaptive Exponential Integrate-and-Fire model [64]

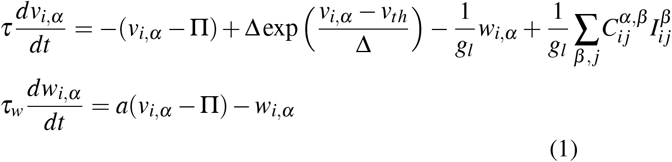

where the sub-threshold dynamics of *i*-th neurons of a certain population *α* (excitatory or inhibitory population of cortical or thalamic network) is described by two coupled state variables (*v_i,α_*(*t*), *w_i,α_*(*t*)), the membrane potential and the adaptation variable, respectively, endowed with discrete reset dynamics. The post-synaptic activity of the *i*-th neuron in population *α* is the collection of emitted spikes {*t_i,α_* } overtime 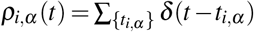. The synaptic current 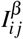 represents the input to the *i*-th neuron given by the activity of *j*-th pre-synaptic neuron belonging to the population *β*. This current is described with a double-exponential conductance-based model

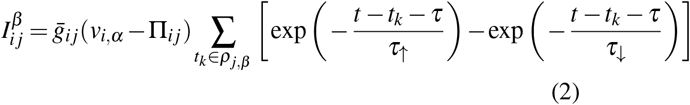

where 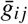 is the maximal conductance, Π_*ij*_ is the synaptic reversal potential, *τ* is the propagation delay and *τ*_↑_ and *τ*_↓_ are the rise and decay time constant, respectively. Every neuron receives a total synaptic current which is the linear sum of such contributions. A coupling matrix *C^αβ^* defines the connections from population *β* to population *α*. All parameter sets have been choosen accordingly with literature and recent experimental findings [17], for further details and parameter values used refer to [14].

Every TC neuron receives an external excitatory input mimicking sensory signals. Every conrtical neuron receives a thalamocortical input and and external excitatory input mimicking ongoing activity from afferent cortical areas. Both external inputs are simulated as Poisson spike trains with different rate parameters. In particular, stimulus unrelated activity from afferent cortical areas is given by different time-varying rates *ν*_0_(*t*) following an Ornstein-Uhlenbeck process

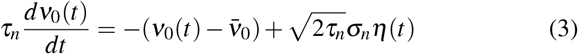

where *η*(*t*) is a Gaussian white noise, *τ_n_* is the characteristic time of stochastic process and *ν*_0_ = 0.75 spk/ms and *σ_n_* = 0.5 spk/ms are the mean and variance, respectively. The parameter values of the stochastic input rate have been chosen in order to match with experimental observations in V1 during external stimulation [65]. Inputs coming from the sensory system are simulated as homogeneous Poisson spike trains with different constant rate values *ν*_ext_ ranging from 0 to 1 spk/ms. We consider two different regimes: one without sensory inputs (*ν*_ext_ = 0 spk/ms) and one with TC neurons receiving external inputs of different rate (*ν*_ext_ > 0 spk/ms). Synaptic currents from external sources are simulated following the same synaptic model as intra-network connections.

### 2.2 Computation of simulated LFP and network firing rate

We define a field quantity related to experimentally recorded mesoscopic LFP signals [72,73,74]. Following [14], we compute cortical LFP_T_(*t*) and thalamic LFP_T_(*t*) as linear combinations of all synaptic intra-network currents. In particular, we sum the absolute values of all synaptic currents to pyramidal (PY) neurons for cortical LFP, and the absolute values of all synaptic currents between TC and RE neurons for thalamic LFP. This simple method of computing LFPs from spiking network models is robust and efficient under the assumption of homogeneous extracellular medium, as shown in [39]. We compute the firing rate of each neural network as the mean of firing rates of all neurons from both excitatory and inhibitory populations

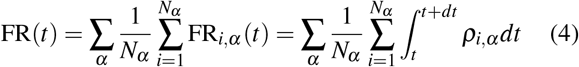

where *α* runs over both excitatory and inhibitory populations. We express thalamic and cortical firing rates in number of spikes per millisecond (spk/ms).

### 2.3 Spectral Analysis

LFP power spectral densities are estimated from a dataset of 40 different simulations, each generated with different noise realisations. We compute the power spectral density (PSD) with a FFT using Welch method *(pwelch* function in MATLAB). To that end, the LFP signal is split up into 8 sub-windows with 50% overlap. The overlapping segments are windowed with a Hamming window. The periodogram is calculated by computing the discrete Fourier Transform, and then calculating the square magnitude of the result. The modified periodograms are then averaged to obtain the power spectral density estimate. We also analyze the distribution of phase lags in frequency-bands of interest, to investigate possible synchronizations of the two networks activities. The cross-spectrum between the thalamic and cortical LFP signals is estimated from the same dataset again using the Welch’s method (*mscohere* function in MATLAB). We also compute the phase coherence as

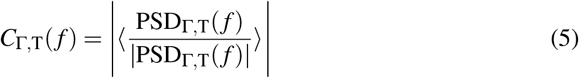

where PSD_T,T_(*f*) is the estimated cross-spectrum and 〈〉 is the average over the dataset. The phase relation between Fourier-transformed signals 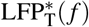 and 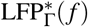 at a given frequency *f* is quantified as the angle of the estimated complex-value cross-spectrum, similarly to [67]. We compute phase-lags △*ϕ_i_*(*f*) for every *i*-th trial in the dataset, and average over frequency-bands of interest (for example the estimated phase-lag for the *δ*-band is the average of phase lags in the 1-4 Hz range). We interpret the estimated phase relations as directional data and we compute the mean resultant vector modulus

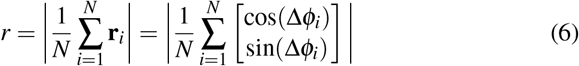

for every frequency-band considered. We test the presence of a preferred phase-lag in a certain frequency band with a Rayleigh test between a uniform distribution hypothesis *H*_0_ and a non-uniform distribution hypothesis *H*_1_, considering the approximated p-value [68]

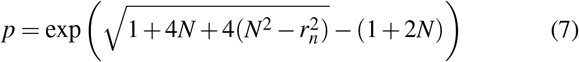

where *N* is the number of sample in dataset and *r_n_* = *rN*. With such a method we quantify if phase-lag values are given by random fluctuations, or whether they belong to a certain non-uniform distribution [69]. We compute filtered Local Field Potential signals in the *δ* and *θ* ranges by means of a digital filter (*designfilt* function in MATLAB). In particular, we consider the first cut-off frequency to be 1/4 Hz and the second cut-off frequency to be 4/12 Hz for the *δ*-range and θ-range, respectively. We use the MATLAB function *filtfilt*, which processes the input data in both the forward and reverse directions to eliminate the phase lag.

### 2.4 Numerical Methods

All scripts are written in MATLAB. The dynamical equations of the system are solved numerically with a second-order Runge-Kutta method with *mid-point* scheme and time step *h* = 0.05 ms. Some scripts have been implemented with the DynaSim Toolbox [66].

## 3 Results

We investigated *in silico* the properties of thalamocortical information transmission through our local network model of the thalamus receiving external stimuli [34] and projecting to the primary cortex receiving stimulus-unrelated excitatory inputs [38] (Figure 1A). We computed the cortical LFP_Γ_(*t*) and the thalamic LFP_T_(*t*) [39] (Figure 1B, see Methods for details) as main outputs of the system.

### 3.1 Frequency-dependent transmission in thalamocortical connections

To capture the mechanisms underlying the relationship between spectral content in the thalamus (both input driven and internally generated) and in the cortex, we first checked whether our model reproduced the lack of spindle oscillations in the cortex experimentally observed [12]. We compared the thalamic PSDT and the associated cortical PSD_Γ_ from the respective LFPs both without external stimuli and during external stimulation, (see Methods and [34]). In the former case both thalamic and cortical networks are associated to prominent *δ* [1-4 Hz] fluctuations, but display several secondary peaks, in the *θ*-band [7-12 Hz] and *γ*-band [30-80 Hz] respectively (Figure 2A). When introducing external inputs, the gamma peak becomes stronger [40] in the cortex, while a secondary beta peak appears in the thalamus (Figure 2B). The corticothalamic ratio, that is the ratio of the cortical and the thalamic spectra across the stimulation range (see Methods), shows that both the *θ*-peak and the *β*-peak are weaker in the cortex (Figure 2B), consistently with experimental observations [12]. This suggests that these bands are specifically suppressed by thalamocortical transmission. We observe that this suppression is poorly modulated by external inputs (Figure 2B). Likewise, external inputs do not affect thalamocortical coupling in slower *δ*-rhythms, which seem to be an intrinsic feature of the system, unrelated to stimuli. External inputs modulation is visible, instead, in the cortical *γ*-band activity, which is increasing proportionally to the input rate (Figure 2B) [41,40,38]. This modulation happens through enhancement of thalamic activity as shown in Figure 2B. The local origin of gamma oscillations in cortical networks is a well known phenomenon [42,43]. Lower cortical frequencies are instead supposed to be phase-locked to thalamic stimuli [44,40,9,38], as is the case in this model (see Figure 3). However, the lack of entrainment of the cortical network to the thalamic inputs between the *δ* and the *γ* bands is a less obvious phenomenon with important consequences for information transmission. In fact, while *δ*-fluctuations are an important component of thalamic activity and they can convey information about the external world during stimulus-driven regimes [45,38] or about the internal brain state during sleep or anesthesia [23,36], *θ*/spindle-oscillations are locally originated in the thalamus [46,47] and hence do not carry information about the external world. Actually, thalamic spindle-rhythms interpose to information transmission and therefore contribute to the gating role of the thalamus [24,48]. Therefore, our results shown in Fig 2 suggest that there are mechanisms in the thalamocortical transmission that implicitly select the informative frequency bands. In the following we will investigate possible mechanisms underlying this selection.

**Fig. 2:**
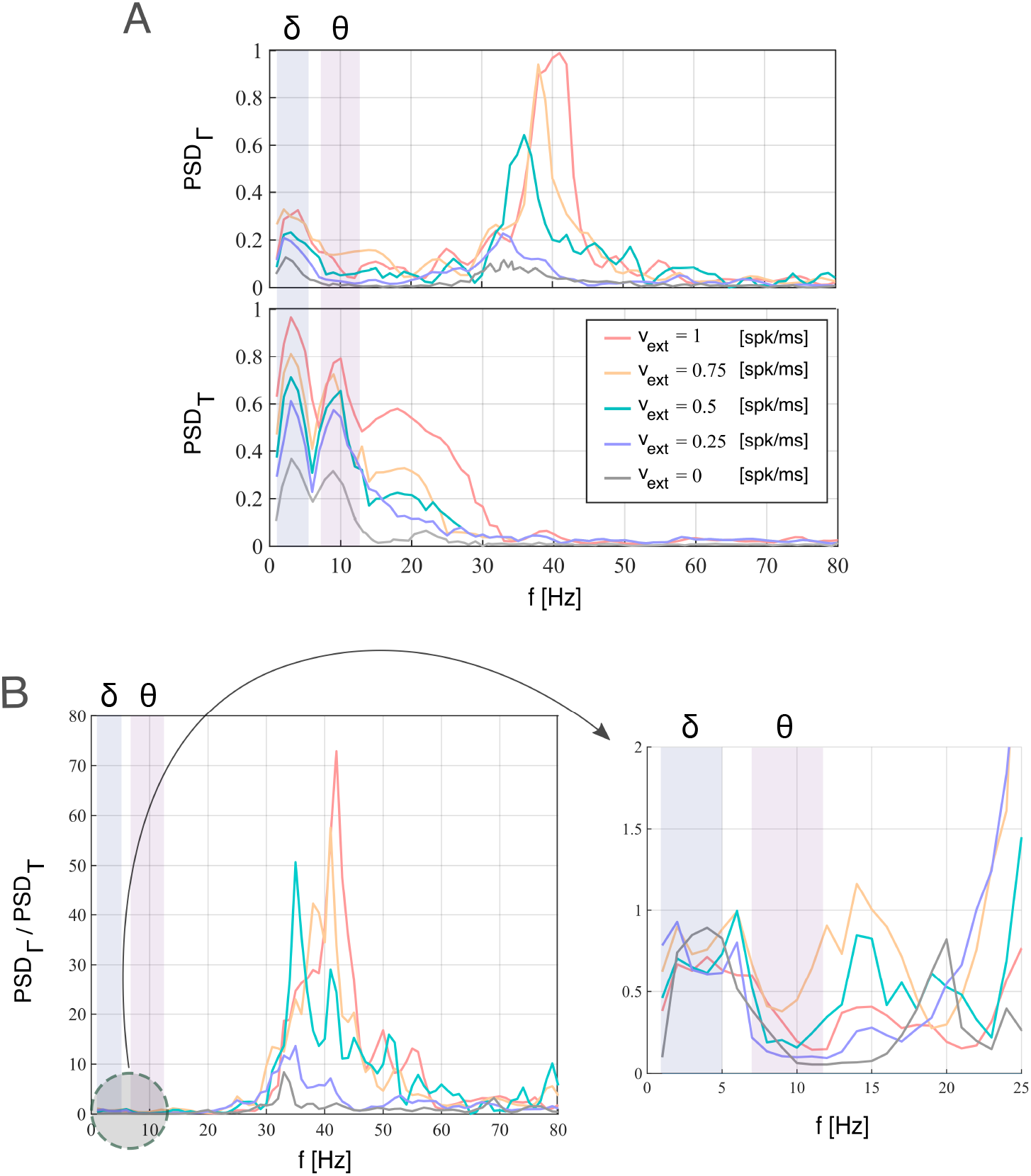
Modulation of thalamocortical LFP spectrum by external input. A) Estimated power spectral densities PSD_T_ and PSD_Γ_ of thalamic and cortical LFP for different external stimulation (the different input intensities are indicated in the legend). The results are averaged over 40 trials and normalized. Each stimulus is simulated as an homogeneous Poisson spike-train with rate *ν*_ext_ (see Methods). The blue and purple stripes represent *δ* [1-4 Hz] and *θ* [7-12 Hz] frequency range, respectively. B) Ratio between cortical and thalamic power spectrum as a function of input rate.

**Fig. 3:**
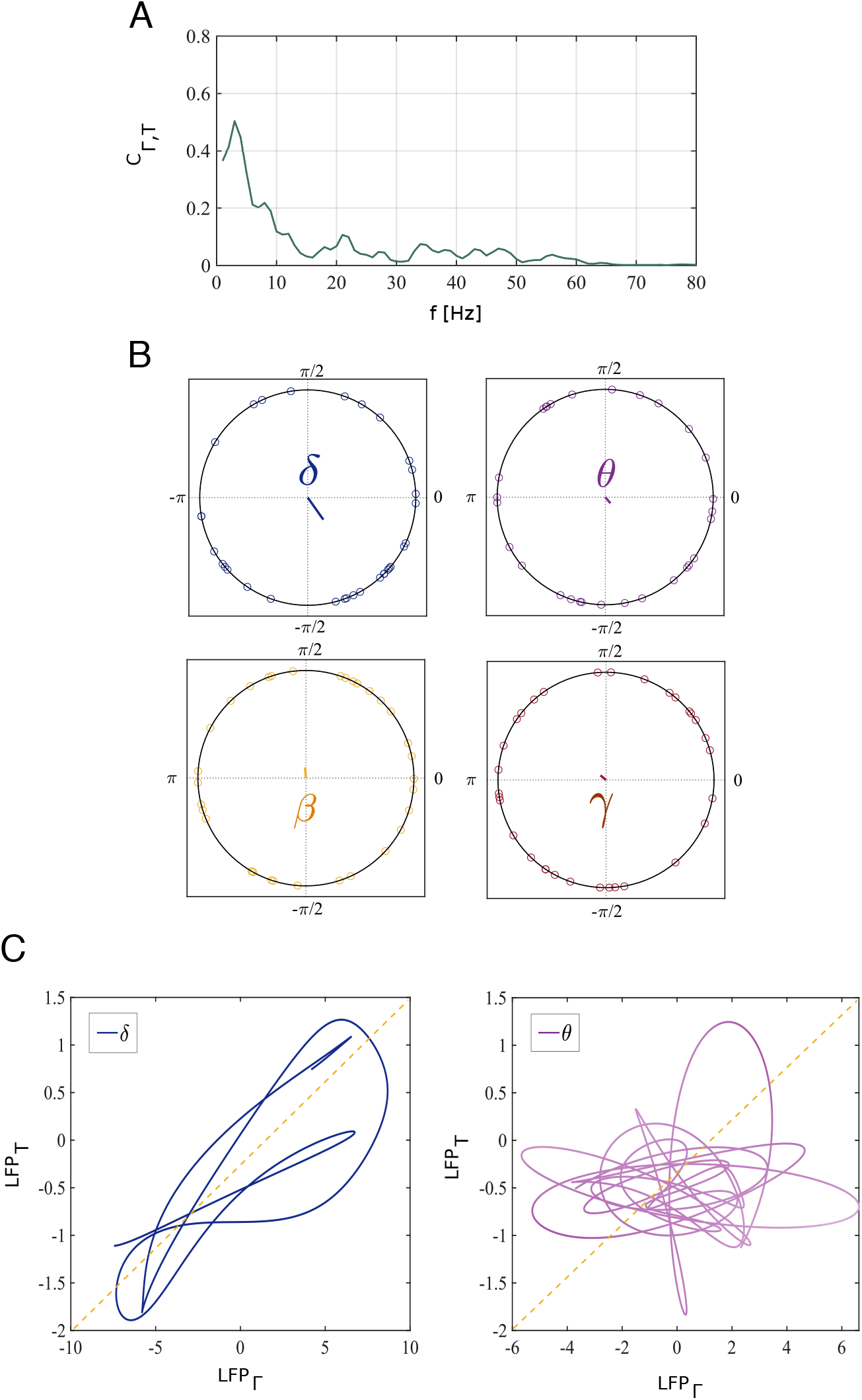
Phase relation between the cortical and thalamic LFPs. A) Phase coherence (see Methods) between thalamic and cortical networks during external stimulation. B) Circular scatterplot of phase-lags between cortical and thalamic LFP signals in four frequency bands: *δ* (1-4 Hz, blue), *θ*/spindle (7-12 Hz, purple), *β* (12-30 Hz, orange) and *γ* (30-80 Hz, red). Empty circles are datasets of 30 different simulations with different external inputs realisations. Lines in the center of the figures are the resultant vector lengths estimated from the dataset. C) Signal trajectory in the (LFP_Γ_, LFP_T_) space for two frequency bands of interest. The blue and purple trajectories represent filtered LFP in the *δ* and *θ* ranges, respectively. Both filtered signals are obtained from the same raw LFP signal for a time-window *T* = 1 s.

### 3.2 Phase relationship between thalamocortical LFP

We asked then whether thalamic inputs regulate cortical activity by influencing its spectral content. We performed a band-wise phase coherence analysis (see Methods), focusing on the frequency bands of interest. We found that *δ* - oscillations are coherent between the thalamic and cortical LFPs (Figure 3A), in agreement with experimental analysis [40], while the system does not show significant phase-coherence in any other frequency bands. Moreover, phase lags in the *δ* band from different simulations tend to cluster around a mean-value, showing that the *δ* oscillations in the cortex are locked to those in the thalamus (Figure 3B). We quantified this observation by computing the resultant vector lengths (see Methods): *R_δ_* = 0.27, *R_θ_* =0.11, *R_β_* =0.08, *R_γ_* = 0.07 (vectors in Figure 3B). We tested the uniformity of the sample distributions with a Rayleigh test (*p*=0.05, see Methods): *p_δ_* < 0.01, *p_θ_* = 0.78, *p_β_* = 0.92, *p_γ_* = 0.93. We then investigated the thalamic and cortical LFP signals filtered in the *δ* and *θ* bands to qualitatively illustrate the different relationship between thalamus and cortex in the two bands (see Methods). The thalamic and cortical LFPs show a certain level of phase coupling in the *δ* range while they behave independently in the *θ* range (Figure 3C). These results show a strong *δ* coupling between the thalamic and cortical networks, while the spectral content in any other frequency band is not coherent within the thalamocortical system. In particular, intermediate *θ* and *β* rhythms in the thalamus do not entrain the corresponding bands in the cortex. We conclude that not only oscillations in the spindle range are much weaker in the cortex than in the thalamus, but the residual oscillations do not have any phase relationship with the spindle oscillations in the thalamus. Therefore hence thalamic spindles are completely filtered out by thalamocortical connections. As before, this suggests that entrainment in *δ* band and filtering of *θ* rhythms is a prominent, stimulus-unrelated characteristic of the thalamocortical system. Such role of slow thalamic rhythms in modulating cortical activity is a recurrent experimental finding [36, 49].

### 3.3 Spindle oscillations in TC neurons

The different behaviors of the frequency bands in thalamocortical transmission necessarily depends on mechanisms at the thalamocortical projection level or inside cortical network dynamics. We focused in understanding these mechanisms by firstly characterizing spindle oscillations in the subset of thalamic neurons projecting to the cortex, rather than in the overall thalamus dynamics. The LFP of a given network is determined by synaptic inputs [72,73] and is simulated in our model accordingly [39]. Consequently, thalamic LFP is mainly composed of synaptic currents of RE inputs into TC neurons and recurrent RE-RE connections, because of their higher connection probabilities respect to TC inputs (see Methods). However, thalamic inputs to the cortex are given only by the output of TC neurons. We compared the spectral features of thalamic LFPT and TC neurons firing at a rate FR_*tc*_ (Figure 4A, see Methods). Specifically, we computed the power spectral density of the thalamic LFP and the TC population firing rate, observing that the *θ* component is much less pronounced in the latter than in the overall LFP. This suggests that part of the discrepancy between thalamic and cortical PSD in the *θ* range [12] is that only a weak component of thalamic spindle oscillations is transmitted to the cortex, as these oscillations are stronger in the RE than in the TC population [34]. Still, even considering the actual input to the cortex, the *δ* band is transmitted in a much more faithful way than the *θ* oscillations (Figure 4B). The ratio between the activities of the overall thalamic network and the TC population only shows increasing discrepancy in the *θ* range, and particularly in the *β* range, for increasing input rate. These discrepancies in the spectral content of network activities rely then on the thalamic connection scheme. The cortical spectrum, in the low frequency range, is indeed much more similar to the cortical input spectrum than to the overall thalamic LFP (Figure 2A). The thalamus modulates cortical activity in the low *δ* rhythms in the absence and in the presence of external sensory stimulation, while processing the sensory input with internal spindle oscillations driven by RE inhibition, in agreement with experiments [36]

**Fig. 4:**
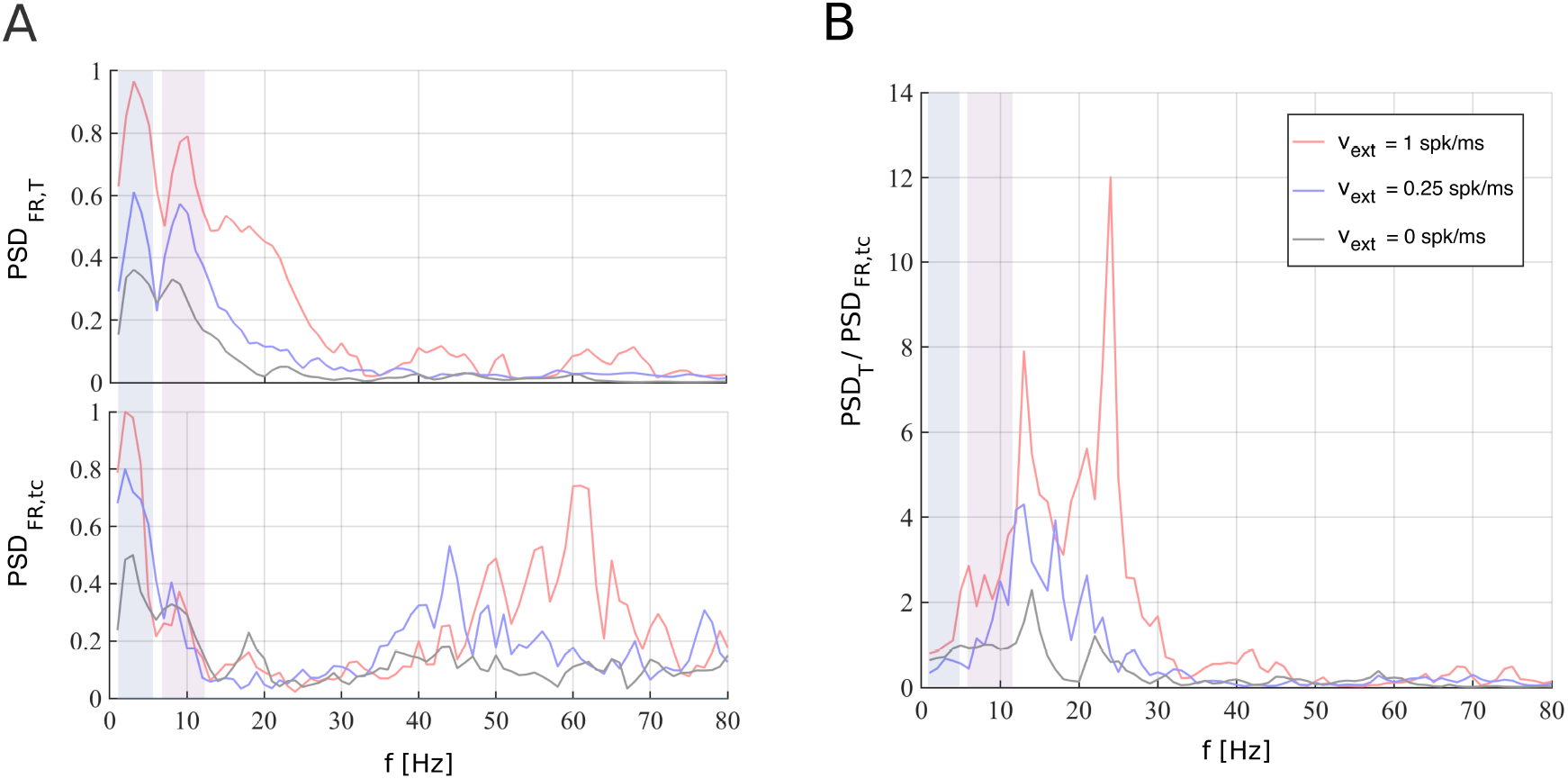
Spectral analysis of thalamic LFP and TC firing rate. Thalamic spectral features in function of different external stimulations (see legend). Blue and purple stripes represent *δ* [1-4 Hz] and *θ* [7-12 Hz] frequency range, respectively. A) Estimated power spectral densities PSD_T_ and PSD_FR,*tc*_ of thalamic LFP (top) and TC firing rate (bottom). Results are averaged over 40 trials and normalized. B) Ratio between thalamic LFP and TC firing rate PSDs.

### 3.4 Modulation of frequency response by internal cortical dynamics

The relatively low component of the thalamic spindle oscillations in the input to the cortex does not account for the whole extent of the filtering of spindle oscillations. We investigated then if the internal cortical dynamics plays a role in shaping the spectral response properties of the cortex as well. To that end, we designed an artificial Poisson spike train input (see Methods) whose rate *ν_m_*(*t*) was composed by the superimposition of three elements: a baseline constant input, a sinusoidal modulation in the δ band, and another one in the θ band, as follows

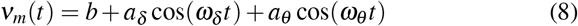

where *b* is the baseline, *a_δ_* and *a_θ_* the amplitudes of the two modulations considered, and *ω_δ_* = 3 Hz and *ω_θ_* = 9 Hz their frequencies. This artificial input is designed to test the sensitivity of the cortical network to specific spectral contents of the thalamic input, and allowed us to analyze the cortical response in function of the ratio Λ between the amplitudes of the two sinusoidal modulations

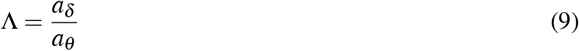

We studied how the cortical spectral response changes with respect to parameter variations in the 2D space (*b*, Λ) (Fig 5A). We found that spectral transmission was symmetric only in the lack of external stimulus, while it favoured *δ* respect to *θ* components for any non-zero value of the external stimulus. For instance, for *b* = 0 and Λ = 0.5 the *θ* component dominates in the cortical activity, while for *b* > 0 the *δ* component is almost as strong as *θ* oscillations, and the opposite does not occur. Overall, the stronger the baseline activity, the higher the peak in the *γ* range and the larger the interval in which the *δ* activity dominates in the cortex, even if injected with a smaller amplitude than *θ*-activity (Fig 5B). These results show that filtering of the *θ* spectral components depends on the cortical sensitivity, and is more severe when the input baseline is strong, and hence the cortical spectrum is dominated by *γ* activity.

**Fig. 5:**
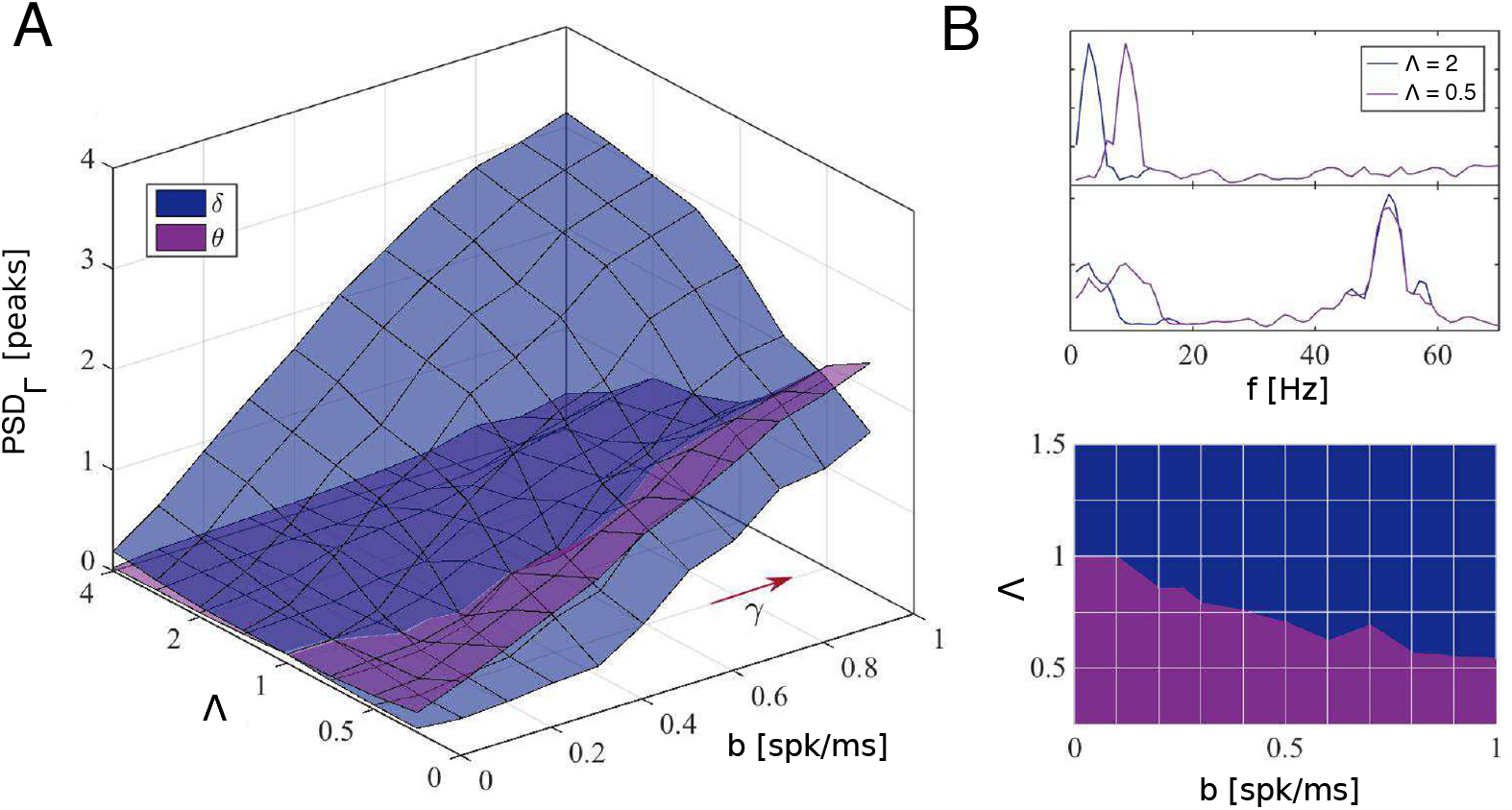
Frequency transfer as a function of network inputs. A) Power peaks in the *δ* (blue surface) and *θ* (purple surface) ranges of estimated cortical PSD when injected with input described in Eq 8 as a function of baseline *b* and ratio Λ between relative amplitudes of the two frequencies of the artificial input. Red arrow represents higher *γ* rhythms increasing in the direction of increasing *b*. The frequency spectrum peaks are in arbitrary units. B) Cortical power spectral density PSD_Γ_ for two extreme values of *b* and Λ (top). Projection on the Λ-*b* plane of intersection between *δ* and *θ* surfaces (bottom). Results are averaged over 20 different simulations. During all simulations we consider *ω_δ_* = 3 Hz and *ω_θ_* = 9 Hz.

### 3.5 Spectral role of the relative weight of thalamic inputs to cortical populations

The results described above reveal that cortical γ rhythms are a key factor for spindle suppression and shifting of information routing within frequency bands. We investigated whether the cortical response depended on ongoing cortical rhythms also during actual thalamic stimulation rather than the simplified external stimulus described in the previous subsection. In particular, we examinated the dependence of the cortical spectrum on the relative weight of thalamocortical inputs to cortical populations. We defined a control parameter *χ^g^* as the ratio between synaptic conductances of thalamocortical afferents 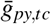 and 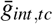 on PY and INT populations, respectively. This latter value was normalised over the ratio 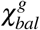 of thalamocortical synaptic conductances considered so far (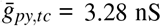 and 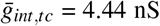, for further details see [14]), obtaining

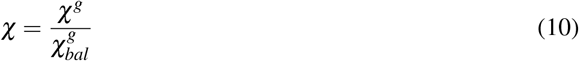

We computed the peaks of cortical PSD_Γ_ in the *δ, θ* and *γ* bands as a function of the control parameter *χ* (Fig 6A). We found that changes in the input balance have a profound effect on the propagation of the different frequency bands from thalamus to cortex observed so far. When INT neurons receive most of the inputs (*χ* < 1), the cortex becomes progressively closer to a faithful reproduction of the thalamic activity (including the *θ* spectral component), up to the disappearance of *γ* oscillations (Fig 6B). When PY neurons receive most of the input (*χ* > 1) the cortex becomes progressively dominated by the *γ* oscillations, due to the interplay with the INT neurons (Fig 6B) disrupting lower frequency oscillations [54]. In the balanced condition (*χ* ~ 1) the *γ* spectral component is present and disrupts the transmission in the *θ* range, but not in the lower *δ* range (Fig 6B). This indicates that a balanced input to the thalamus is the optimal way to selectively block only the non-informative *θ* component generated by the thalamic internal activity [23, 55], so that *γ* rhythms are thus the decisive factor of this mechanism.

**Fig. 6:**
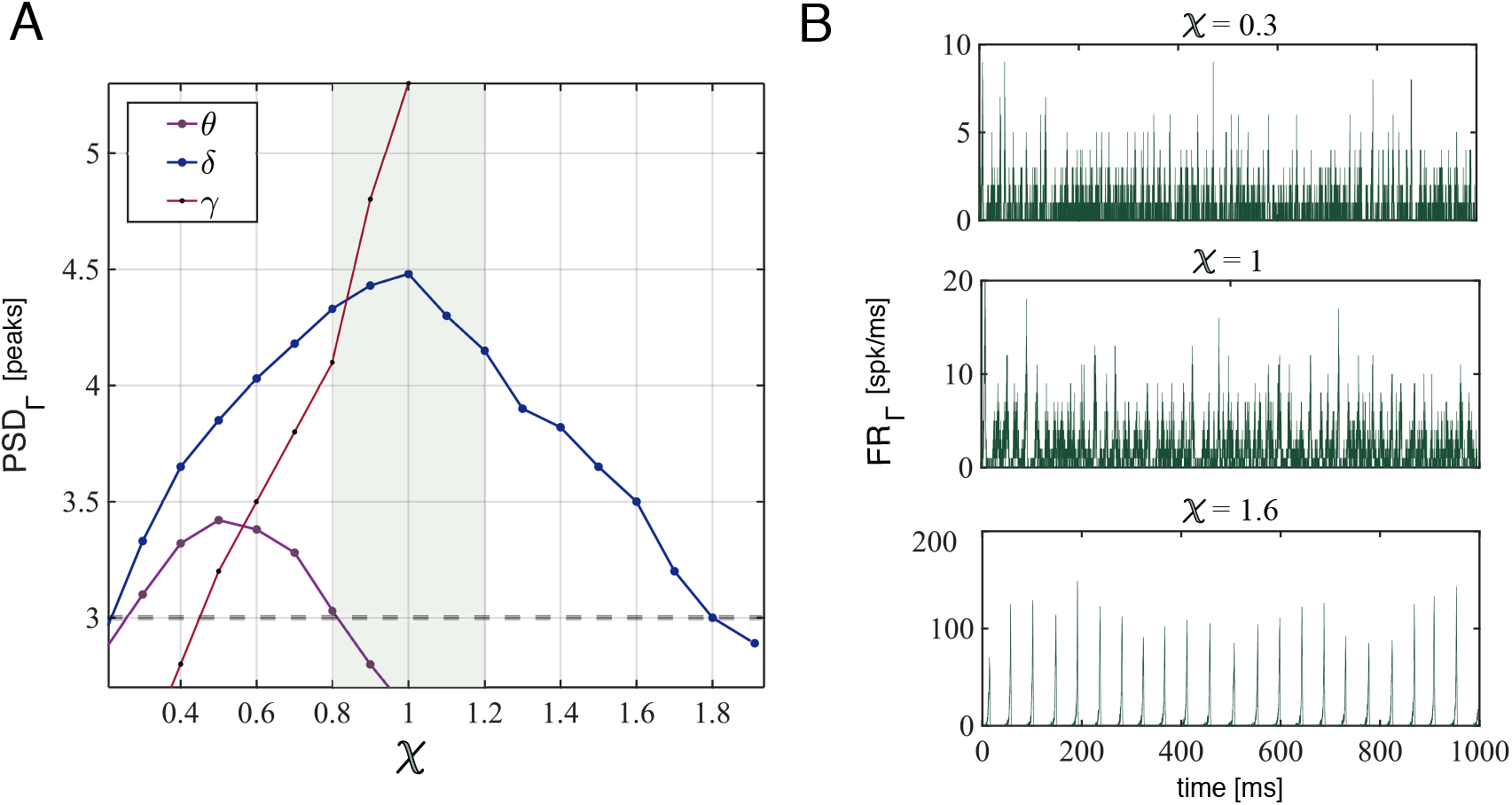
Spectral cortical and thalamocortical architecture. A) Modulation of the cortical PSD_Γ_ as a function of control parameter *χ*. Line with markers indicate power peaks for different frequency bands (see legend). Light green indicates the reference region. Power under the grey dashed line are not associated to peaks. The spectrum peaks are in arbitrary units. B) Firing activity of cortical network for different values of *χ* (from top to bottom: *χ* = 0.3, 1, 1.6). Results are averaged over 20 trials.

## 4 Discussion

Our *in-silico* investigation shows that thalamocortical connectivity modulates spectral transmission between the two areas in a way that optimizes spectral information transmission. We observed that thalamic oscillations in the *δ* band [1-4 Hz] and in the θ/spindle band [7-12 Hz] have remarkably different effects on cortical activity. In fact, the latter embodies slower δ with respect to thalamic spindles rhythms while showing another frequency-peak in the higher *γ* band [30-80 Hz]. Under external stimulation, the cortex reproduces the thalamic *δ* band and locally generates *γ* rhythms proportionally to the input intensity, while the thalamic spectra show an enhancement in the *β* band [20-30 Hz]. This model reproduces well-known cortical *γ* peaks in response to increasing external inputs [42,43], but it also differentiates functional roles of thalamic spectral components in transmitting information to the cortex. In particular, it suggests that thalamic *δ* rhythms act as a local *clock* conveying information about internal thalamic states through the activity of the reticular nucleus, in agreement with experimental findings [36]. Our results suggest that external stimulation modulates high-frequency activities over a low-frequency background activity, keeping δ rhythms as a communication channel, similarly to what happens during non-REM sleep [56]. On the other hand, thalamic spindles in the *θ* band play a minor role in communication to the cortex [55], as they are locally generated in the thalamus. By investigating the mechanisms behind spectral selectivity, we found that both thalamocortical afferents and cortical responsiveness play a role in shaping spectral information transmission, as we dissect in what follows.

Firstly, thalamocortical afferents are composed only by projections from thalamocortical relay (TC) neurons to cortical populations. However, the thalamic LFP is mainly contributed by reticular (RE) projections, which embody stronger *θ* rhythms [34] arising from an inhibition-driven interplay between RE and TC cells, as observed experimentally [23, 25]. Consequently, the spectral composition of thalamocortical inputs reveals that only a fraction of *θ* rhythms is really conveyed to cortical populations. Our results strengthen the hypothesis that spindles, commonly observed in the thalamus [23] are a localized feature of the thalamic network which serve as modulators of information trasmission from sensory input to the cortex through tuning of thalamocortical relay modes [24]. Indeed, spindles are locally generated in the thalamus and typically arise from the interplay between TC and RE neurons [13]. In particular, RE neurons shape the computational capabilities of the thalamus through the low-threshold bursting behavior of TC neurons. Reticular neurons hyperpolarize TC neurons leading to de-inactivation of T-type calcium channels, what switches TC activity from tonic to bursting [24], leading to spindle oscillations. Many attempts have been made in order to reproduce spindle activity in computational models of thalamic circuitry. One of the first works on TC-RE was developed more than 20 years ago [29]. Recent works describe spike-wave discharges in sleeping thalamus by means of neural field models [30]. Other late works are based on modeling thalamic cells using single-compartment conductance-based model with several ionic currents. One example is [31], in which the authors suggest that intrinsic thalamic activities are able to generate oscillations at different timescales, from slow waves to gamma rhythms. However, state-of-the-art modelling of thalamic oscillations lacks simpler yet efficient integrate-and-fire network models. Apart from our work (related to [34]) we are only aware of a single previous work regarding Up-Down oscillatory activity in thalamus and cortex [33].

Secondly, we characterized an intrinsic mechanisms of frequency selection as a function of the cortical dynamical state. Previous works report that cortical *γ* oscillations respond in a resonant manner, implying the proximity of cortical dynamics to a Hopf-bifurcation [57, 58]. We extend this by showing that the absence/presence of *γ* rhythms shape the responsiveness of cortical network with respect to external modulation in the *δ* and the *θ* band. In particular, the presence of strong stimulus-driven *γ* oscillations seems to dampen cortical resonance in the *θ* range while acting as a slow envelope in the *δ* range. When *γ* rhythms are absent, the cortical network behaves in the asynchronous irregular (AI) [43] state and thus shows a symmetrical frequency-response portrait.

Finally, we also investigated the role of cortical synaptic parameters in shaping the cortical response, focusing on thalamocortical synaptic conductances. We showed that, in our modeling description, a kind of *democratic* thalamocortical relay to both cortical population through TC neurons acitivyt is optimal for information routing. This input balancing reflects the balanced-network approach used in modeling cortical networks, which is able to reproduce sub- and supra-threshold fluctuations similar to recordings *in vivo* [59, 60], and is thus a promising theoretical framework to study cognitive processes in the cortex [61, 62]. In fact, cortical γ rhythms arise from such a balance scheme as a collective phenomenon resulting from the interplay between excitation and inhibition [42]. In our case, balanced inputs correspond to thalamocortical afferents sending stronger inputs to interneurons respect to pyramidal neurons in order to maintain their internal balance. Such scheme is in agreement with recent experimental observations [17, 16]. Seminal works by Sherman such as [15] highlight that thalamocortical inputs to the cortex are characterized by large and sparse connectivity, able to send powerful information throughout the highly interconnected cortical circuitries. Moreover, thalamocortical relays seem to account for 5-25% of the excitatory connections within the cortex [16]. *In vivo* recordings suggest that thalamocortical afferents seem to target in equal numbers to both the cortical excitatory and inhibitory populations [17]. To the best of our knowledge, our work is the first effort in modelling the mesoscopic activity arising from such a sparse but symmetric architecture by means of a simple spiking neural network.

Investigations of thalamocortical activity have been done by Bazhenov et al [18] with a model based on *in vivo* recordings, which was able to reproduce Up-Down oscillations. However, the authors considered few neurons with complex Hodgkin-Huxley models and fixed connectivity scheme. Later, Izhikevich and Edelman [19] developed a large-scale model of remarkable complexity with several cortical layers, synaptic plasticities and anatomical features. This model is based on the powerful-and-sparse approach for thalamocortical feedforward afferents and on a simple (integrate-and-fire) Izhikevich model for single neuron dynamics [20]. Recently, higher levels of detail have been reached in modeling thalamic and cortical oscillations [21] and local field potentials with 3D distribution models [22]. Despite of these relevant works, a simple and manageable model able to test novel experimental findings such as [16,17] is a necessary step to have a shot at describing thalamocortical dynamics while easily investigating different parameter settings.

Computing local field quantities from the activity of spiking network models gives a wider view of the relationship between neuronal interactions and collective phenomena such as oscillations and synchronization [63]. In this respect, it would be of interest to test experimentally the relationship between cortical balance and spectral content transmission by means of cortical stimulations and LFP recordings. As we focused on particular thalamic input emerging from our thalamic model [34], a possible further development would be to design different artificial inputs and analyze the corresponding cortical response as a function of both stimuli features and the internal dynamical state of the network. We stress that here we have focused on feedforward interactions from a local thalamic area to a primary cortical circuitry of the related sensory system. Our model prediction can be used, for instance, to study the effects of sensory inputs coming from the retina to a local circuitry of the LGN, conveying information to a local circuitry of the primary visual cortex V1. A possible extension of this work would be adding cortical feedback to the thalamocortical system and investigate novel collective organisations of the circuit in response to external stimulation.

## Acknowledgements

J.G.O. was supported by the Spanish Ministry of Science, Innovation and Universities and FEDER (project PGC2018-101251-B-I00 and Maria de Maeztu Programme, project CEX2018-000792-M), the Generalitat de Catalunya (project 2017 SGR 1054), and the ICREA Academia pro- gramme.

E.C. was supported by the grant “PANACEE” (Prevision and analysis of brain activity in transitions: epilepsy and sleep) of the Regione Toscana – PAR FAS 2007–2013 1.1.a.1.1.2 – B22I14000770002.

AM was supported by the Italian Ministry of Research (MIUR, PRIN2017, PROTECTION, project 20178L7WRS).

